# Signal complexity indicators of health status in clinical-EEG

**DOI:** 10.1101/2021.03.16.435656

**Authors:** Kelly Shen, Alison McFadden, Anthony R. McIntosh

**Affiliations:** Rotman Research Institute, Baycrest Centre; University of Toronto

**Author notes:** Corresponding Author, Mailing Address: 3560 Bathurst Street, Toronto ON M6A 2E1, Canada.

**Keywords:** Multi-scale entropy, EEG, neurodegenerative disease, epilepsy

## Abstract

Brain signal variability changes across the lifespan in both health and disease, likely reflecting changes in information processing capacity related to development, aging and neurological disorders. While signal complexity, and multiscale entropy (MSE) in particular, has been proposed as a biomarker for neurological disorders, most observations of altered signal complexity have come from studies comparing patients with few to no comorbidities against healthy controls. In this study, we examined whether MSE of brain signals was distinguishable across individuals in a large and heterogeneous set of clinical-EEG data. Using a multivariate analysis, we found unique timescale-dependent differences in MSE across various neurological disorders. We also found MSE to differentiate individuals with non-brain comorbidities, suggesting that MSE is sensitive to brain signal changes brought about by metabolic and other non-brain disorders. Such changes were not detectable in the spectral power density of brain signals. Our findings suggest that brain signal complexity may offer complementary information to spectral power about an individual’s health status and is a promising avenue for clinical biomarker development.

## Introduction

A growing literature suggests that some degree of brain signal variability is vital to optimal brain function. Although seemingly paradoxical, noisy (or complex) brain signals are related to a greater capacity for information processing as compared to more predictable signals (Garrett et al., 2018; Vakorin and McIntosh, 2012). Sample entropy is one way to capture the variability of a brain signal (Richman and Moorman, 2000) and multiscale entropy (MSE), where complexity is examined across multiple timescales (Costa et al., 2005), has been particularly useful in broadening our understanding of the role of noise in brain health and disease. MSE, like other measures of entropy, captures the variability in a signal but can additionally differentiate variability induced by increasing randomness, such that white noise gives lower MSE values (Costa et al., 2005). An increase in MSE has been observed in tasks requiring memory retrieval (Heisz et al., 2012) or the integration of stimulus features (Misić et al., 2010) and seems to support accurate and stable behavior (Misić et al., 2010; Raja Beharelle et al., 2012). MSE has been shown to have timescale-dependent shifts during brain development (Hasegawa et al., 2018; Lippé et al., 2009; Miskovic et al., 2016; Szostakiwskyj et al., 2017) and aging (McIntosh et al., 2014; Sleimen-Malkoun et al., 2015; H. Wang et al., 2016) that supports cognitive function (Heisz et al., 2015; Yang et al., 2013), reflecting changes in the brain’s information processing capacity across the lifespan. MSE also reflects processing capacity changes related to various brain diseases including dementia (Bertrand et al., 2016; Grieder et al., 2018; Niu et al., 2018), neurodevelopmental disorders (Mišić et al., 2015; Takahashi et al., 2016; Weng et al., 2017), and psychiatric disorders (Hager et al., 2017; Takahashi, 2013; Yang et al., 2015).

In nearly all of these studies, brain signal complexity changes related to various brain diseases have been detected by comparing individuals with few to no comorbidities against matched healthy controls using data collected in highly controlled laboratory environments. While MSE has been proposed for use as a clinical biomarker for various neurological disorders (Jeste et al., 2015; Lu et al., 2015; Tsai et al., 2015), whether differences in brain signal complexity can be detected across individuals of a heterogenous clinical population per se remains unknown. In this study, we leveraged the Temple University Corpus EEG database (Obeid and Picone, 2016) to test the utility of MSE as an indicator of health status in a large and heterogeneous clinical population. We found MSE of clinical-EEG signals differentiated individuals of varying brain disorders. Interestingly, we also found MSE to differentiate between individuals with non-brain comorbidities and those without comorbidities.

## Methods

### Subjects

Clinical EEG data and corresponding physician reports were downloaded from the Temple University Hospital EEG Epilepsy Corpus (v0.0.1) containing 100 subjects deemed to have epilepsy and 100 subjects without epilepsy (https://www.isip.piconepress.com/projects/tuh_eeg/) (Obeid and Picone, 2016). Subjects from the epilepsy group were included in our sample if the report indicated a previous diagnosis of epilepsy, if the EEG supported a diagnosis of epilepsy, or if the patient had experienced 2 or more unprovoked seizures occurring more than 24 hours apart and the EEG did not contraindicate epilepsy. Subjects without epilepsy were included if they did not meet any of these criteria. Subjects from either group were excluded if a seizure occurred during the recording, if the subject’s level of consciousness was decreased, or if the subject was under the effect of a device likely to cause substantial EEG artifact such as a pacemaker or ventilator. Subjects were also excluded if their recordings were deemed unsuitable in the preprocessing stage due to the presence of artifacts.

Demographic and clinical characteristics were extracted from the physician reports (Table 1). For the various brain-acting medications (anti-epileptic drugs, barbiturates, benzodiazepines, antipsychotics, and antidepressants), subjects were considered to be on them if their medication list included at least one medication of that category. The total number of other (i.e., not brain-acting) medications for each subject was computed by counting the number of total medications listed for the subject and subtracting the number of medications that fell into the brain-acting medication categories listed above. If the medication list stated “others” or a pluralized general category of medications (i.e. “antihypertensives”), two medications were added to the non-brain medication count. Most of the non-brain acting medications reported (69.3%; 223/322) are those used to treat cardiovascular disease, diabetes or chronic respiratory illness. Seizure classifications and terms were determined as outlined by the International League Against Epilepsy (Berg et al., 2010; Blume et al., 2001). A subject was considered to have experienced generalized or focal seizures if their physician’s report contained either a diagnosis falling in one of those categories or a description of seizures matching the expected presentation for that seizure classification. Thirty-four subjects experienced seizures of unknown classification and were excluded from analysis. A further 3 subjects did not have age or sex information available and were also excluded from analysis. This resulted in a total sample size of 163 subjects. Accepted phrases for stroke included indication of a past or present ischemic stroke, hemorrhagic stroke, “CVA”, or intracerebral bleed. Accepted diagnoses for degenerative brain diseases included Alzheimer’s disease, Parkinson’s disease, and dementia. Accepted diagnoses for psychiatric disorders included anxiety, depression, bipolar disease, and schizophrenia. Accepted diagnoses for neurodevelopmental disorders included Down’s syndrome, ADHD, intellectual disabilities, and cerebral palsy. Finally, other brain disorders and injuries included head trauma, brain surgery, brain cancer or metastases, hypoxic brain injuries, encephalitis and meningitis.

**Table 1.** Demographic and clinical characteristics of study sample.

| *Variables* | *Subjects*<br>*(n=163)* |
| --- | --- |
| Age, mean (SD, range) | 52.12 (19.88, 7-91) |
| Sex, *n* female (%) | 91 (55.83) |
| *Medication use* |  |
| Anti-epileptic drug use, % | 36.2 |
| Barbiturate use, % | 2.45 |
| Benzodiazepine use, % | 11.04 |
| Antipsychotic use, % | 9.82 |
| Antidepressant use, % | 11.04 |
| Other medications, mean (SD, range) | 2.26 (2.75, 0-13) |
| *Past medical history* |  |
| Diagnosis of epilepsy, % | 40.49 |
| History of stroke, % | 19.02 |
| Diagnosed degenerative brain disease, % | 4.91 |
| Diagnosed psychiatric disorder, % | 12.27 |
| Diagnosed neurodevelopmental disorder, % | 3.68 |
| Other brain disorder or injury, % | 16.56 |

### EEG preprocessing & analysis

Each subject contributed one EEG recording. For subjects with multiple recordings, the recording corresponding to the physician report containing the most complete clinical picture was selected. For recordings that were split into multiple segments, the longest of the segments was chosen for preprocessing. All preprocessing was performed using the FieldTrip toolbox in MATLAB (www.fieldtriptoolbox.org) (Oostenveld et al., 2011). For each selected recording, 19 scalp electrodes of the International 10-20 system that were common to all subjects were selected. The resulting continuous recordings were segmented into 4-s trials, producing an average of 317 trials per subject, and bandpass filtered (0.5 to 55 Hz). The majority of recordings were sampled at 250 Hz, but one subject that was sampled at 512 Hz was downsampled to 250 Hz before proceeding.

Two trial removal steps were then completed. The majority of subjects received photic stimulation. For these subjects, trials where photic stimulation began and ended were detected, and the trials within this range to 5 trials past the end of stimulation were removed. Trials at the beginning of a recording where the amplitude of the photic channel was not zero were also removed. Next, trials with excessive signal amplitude were detected for removal. For each subject, 30% of the trials that were determined by visual inspection to be reasonably free of artifacts were selected. Global field power was calculated and its mean ± 5 std was used to reject trials with time points outside of this threshold. The average number of remaining trials per subject following both of these removal steps was 178.

Independent component analysis was next used to remove ocular and muscle artifacts. Components with topographical distributions typical of these artifacts were selected and their traces further examined. Where possible, probable ocular artifact components were confirmed via alignment of the component trace with the electrooculogram traces from the original recording. Probable muscle artifact components were confirmed by the presence of a high frequency component trace. Finally, any recordings not referenced to a common average were re-referenced.

MSE (Costa et al., 2005, 2002) was computed by first coarse-graining the EEG time series of each trial into 20 scales. To produce the time series coinciding with a given scale *t*,data points from the original time series within non-overlapping windows of length *t* were averaged. Thus scale 1 represents the original time series, with 1000 data points per channel per trial resulting from 4 seconds of recording sampled at 250 Hz. Next, sample entropy was calculated for each time series across all scales. This measured the predictability of the amplitude between two versus three consecutive data points (m=2), with the condition that data points were considered to have indistinguishable amplitude from one another if the absolute difference in amplitude between them was ≤50% of the standard deviation of the time series (r = 0.5). The resulting values were averaged across trials to produce a single MSE curve per channel for each subject. As an entropy-based measure, MSE values are low for both completely deterministic as well as completely uncorrelated signals.

Changes in MSE occur with changes in spectral power (Lippé et al., 2009; McIntosh et al., 2008) so we additionally assessed spectral power (SPD) alongside MSE. SPD was calculated for each trial using the fast Fourier transform with a Hann window. To account for age-related global signal power changes, each recording was first normalized (mean = 0, *SD* = 1). Relative spectral power was then calculated for each trial, and results averaged across trials to acquire mean SPD per channel for each subject.

### Partial Least Squares Analysis

MSE and SPD measures were each correlated with the available demographic and clinical data using a Partial Least Squares (PLS) analysis (Krishnan et al., 2011; McIntosh and Lobaugh, 2004). This multivariate statistical approach identifies a set of latent variables (LVs) that represent the maximal covariance between two datasets. First, the correlation between the MSE/SPD and clinical data was computed across subjects. Singular value decomposition was then performed on the correlation matrix to produce LVs, each containing three elements: 1) a set of weighted “saliences” that describe a spatiotemporal brain pattern of MSE/SPD measures; 2) a scalar singular value that expresses the strength of the covariance; and 3) a design contrast of correlation coefficients that express how the clinical data relate to the saliences. The mutually orthogonal LVs are extracted in order of magnitude, whereby the first LV explains the most covariance between MSE/SPD and clinical data, the second LV the second most, and so forth. The significance of each LV was assessed with permutation testing by randomly reordering subjects’ MSE/SPD pairing with clinical data to produce 1000 permuted sets for singular value decomposition, with the set of 1000 singular values forming the null distribution. The reliability of the MSE/SPD at each electrode in expressing the covariance pattern of each LV was assessed using bootstrap resampling. A set of 500 bootstrap samples was created by resampling subjects with replacement. The ratio between the saliences and the estimated standard error (bootstrap ratio) was taken as an index of reliability. With the assumption that the bootstrap distribution is normal, the bootstrap ratio is akin to a Z-score and corresponding saliences are considered to be reliable if the absolute value of their bootstrap ratio is >= 2. For the clinical data, confidence intervals were calculate from the upper and lower bounds of the 95^th^ percentile of the bootstrap distribution of the correlation with the scores from the MSE/SPD data. The scores are the dot-product of the saliences with the data for each subject and are similar to a factor score from factor analysis.

For the demographic and clinical data entered into the PLS analysis, age and number of non-brain medications were treated as continuous variables, while all other variables were categorical. Sex was coded as 0 (F) and 1 (M). The remaining variables were coded as 0 (not on drug or does not have condition) or 1 (on drug or has condition).

## Results

To determine whether different and heterogeneous clinical profiles can result in differences in brain signal complexity, MSE curves for each subject were correlated with their demographic and clinical data using a PLS analysis. The singular value decomposition of the correlation matrix resulted in two significant LVs. The first LV showed a differentiation between brain disorders, with a global shift towards greater signal complexity in finer time scales and lower signal complexity in coarser time scales across all electrodes for subjects who experienced generalized seizures or those taking antidepressants as compared to those with other brain conditions (i.e., focal seizures, stroke, neurodevelopmental disorders) or using other medications (i.e., anti-epileptics, barbiturates) (Fig. 1A-B). This shift in MSE was evident when a median-split was performed to classify subjects according to how much they expressed the patterns of the LV (i.e., a median split of the LV-scores, Fig. 1C). This LV was significant (p < 0.001) and accounted for 57.8% of the covariance in the data.

**Figure 1.**
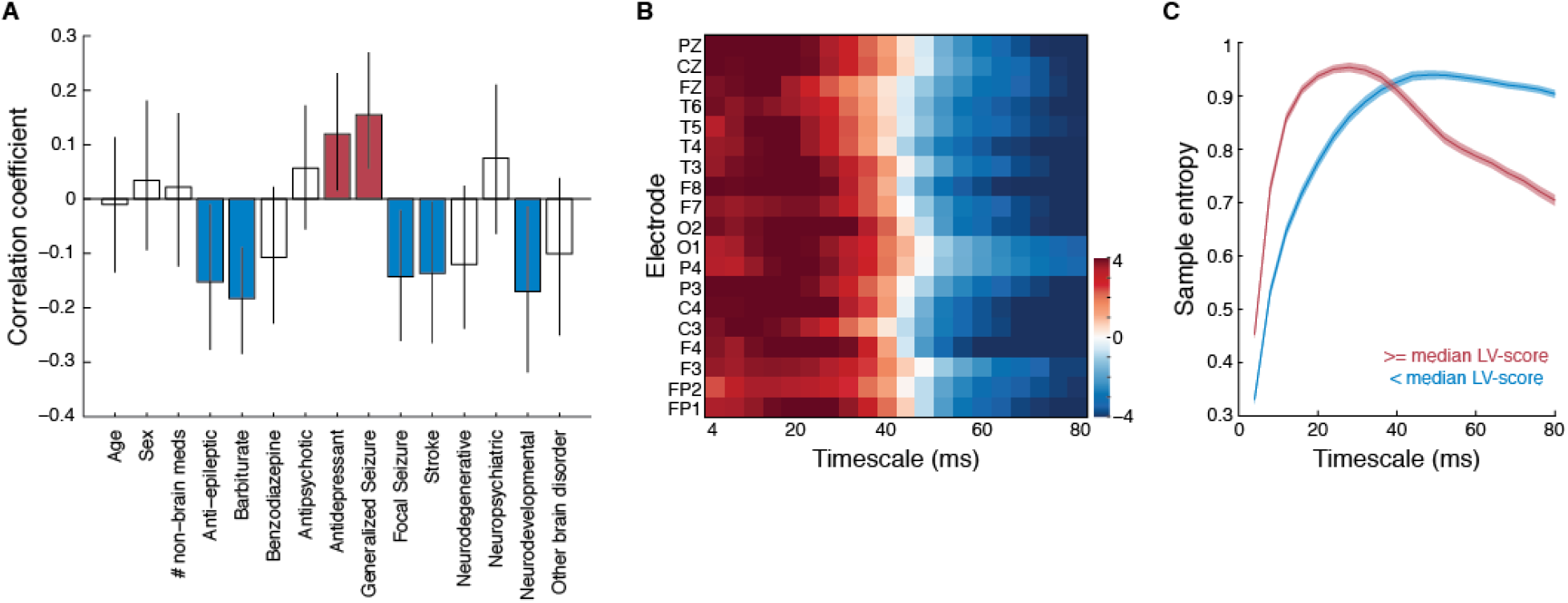
Brain signal complexity differentiates brain disorders. (A) Correlation coefficients and (B) bootstrap ratios of the first latent variable relating clinical data to MSE curves. (C) Average (± SEM) MSE curves, with subjects split into two groups according to their LV-scores. MSE curves were first averaged across electrodes within subjects, then averaged across subjects within each group. In (A), variables whose coefficients are significantly different from 0 are indicated in color for ease of interpretation.

The second LV differentiated older unhealthy (as indexed by the number of non-brain-related medications taken) males who did not have neurodegenerative disease from other subjects, and was associated with slightly higher entropy at the very finest scales and lower brain signal complexity across more coarse time scales (Fig. 2). This LV was significant (p < 0.01) and accounted for 29.0% of the covariance in the data.

**Figure 2.**
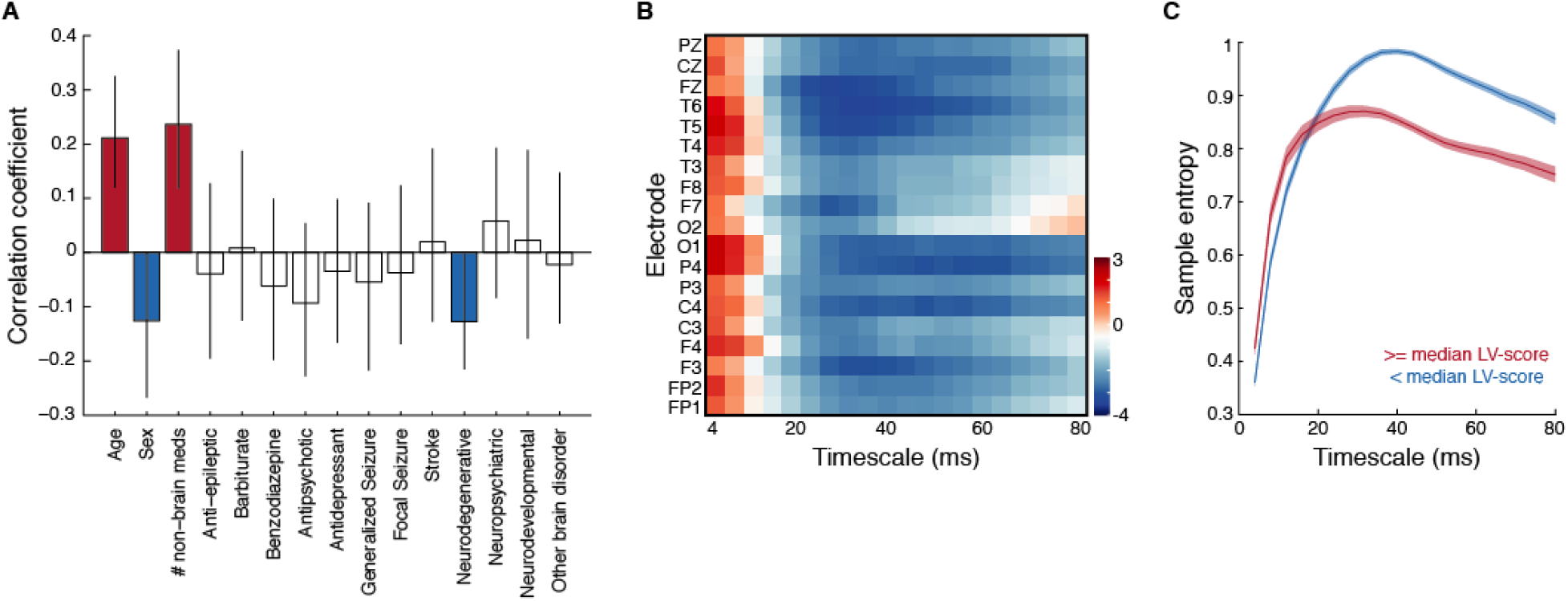
Brain signal complexity differs for older unhealthy males. Correlation coefficients (A) and bootstrap ratios (B) of the second latent variable relating clinical data to MSE curves. (C) Average (± SEM) MSE curves, with subjects split into two groups according to their LV-scores. MSE curves were first averaged across electrodes within subjects, then averaged across subjects within each group. In (A), variables whose coefficients are significantly different from 0 are indicated in color for ease of interpretation.

The MSE profiles for each of the latent variables therefore reflected a unique timescale-dependent shift in brain signal complexity associated with different brain and non-brain disorders. A similar analysis of SPD indicated that SPD profiles could only differentiate between subjects with epilepsy from those without epilepsy (Figure 3).

**Figure 3.**
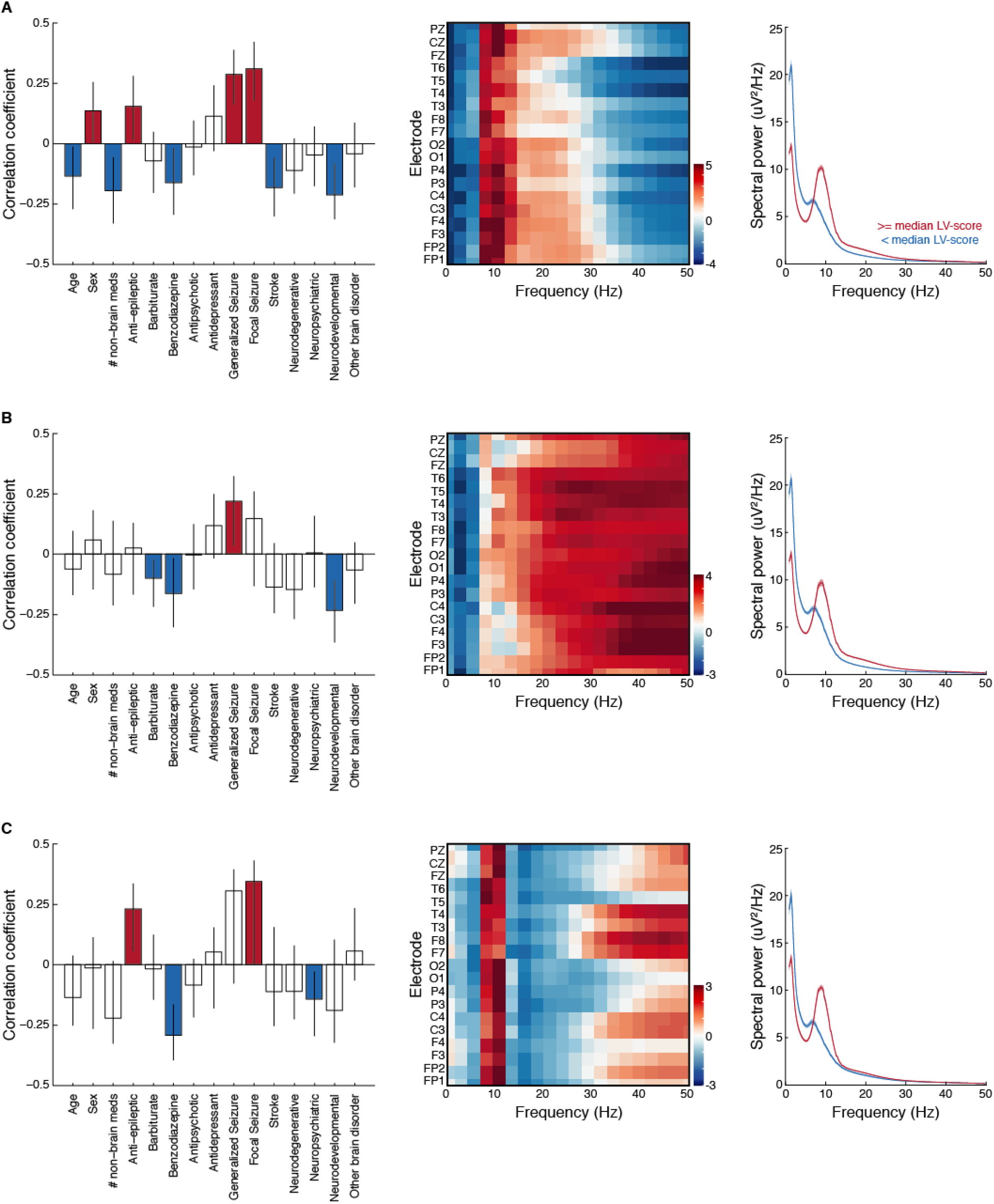
Spectral power density differentiates epilepsy from other brain disorders. (A) First latent variable (p <0.001; 50.1% covariance explained) (B) second latent variable (p < 0.001; 29.9% covariance explained), and (C) third latent variable (p = 0.068; 10.1% covariance explained) of a PLS analysis relating clinical data to SPD. Left panels: correlation coefficients; middle panels: bootstrap ratios; right panels: average (± SEM) SPD, with subjects split into two groups according to their LV-scores. SPD functions were first averaged across electrodes within subjects, then averaged across subjects within each group.

## Discussion

In this study, we examined whether brain signal complexity varied across individuals of a large and heterogeneous clinical population using a data driven approach. We found timescale-dependent differences in brain signal complexity for individuals who experience generalized seizures from individuals who have other brain disorders (e.g., focal seizures, stroke, neurodevelopmental disorders). We also found a timescale-dependent shift in brain signal complexity for older males on various medications not related to neurological or neurodegenerative disease that was not evident in the spectral power of the clinical-EEG recordings. Our findings suggest that brain signal complexity, as indexed by MSE, can provide additional insights into brain health status and function not captured by spectral power.

In line with the notion that the brain is a dynamical system in which “noise” allows for flexible functioning and a variety of metastable states (Deco et al., 2017, 2011), MSE can be considered as an index of functional repertoire (Heisz et al., 2012). Changes to brain function and dynamics can occur with neurological disease and, indeed, differences in MSE from matched controls have been reported for both epilepsy (Weng et al., 2015) and neurodegenerative disease (Tsai et al., 2015). Here we build on these previous reports by showing how the changes in MSE in these neurological conditions can be differentiated from each other. A complementary data-driven analysis of SPD showed changes in power across frequency bands that differentiated epilepsy from all other diagnoses as well as generalized from focal seizures, consistent with numerous accounts of SPD differences in epilepsy (Clemens et al., 2000; Díaz et al., 1998; Niso et al., 2015; Quraan et al., 2013; Walker, 2008). However, the differentiation of individuals with non-neurological comorbidities was unique to MSE.

The MSE results also replicate previous observations that the scale-dependent changes are indicative of neurodegenerative disorders (Figure 2). Higher MSE at coarse-scales was shown to predict cognitive decline in Parkinson’s patients who would develop dementia (Bertrand et al., 2016). The relative balance within subjects between fine and coarse scales also relates to cognitive status in aging (Heisz et al., 2015). These results, considered in the context of the present data, suggest that the relative shifts of complexity across temporal scales may be a sensitive index to assist in clinical evaluation, particular as a predictor of future cognitive decline (McIntosh, 2019).

Metabolic diseases such as diabetes mellitus are known to affect brain structure and cognitive function (Soininen et al., 1992; Tan et al., 2011). More recently, changes to resting-state functional networks have been observed in individuals with diabetes mellitus compared to controls (Y. F. Wang et al., 2016). Autonomic dysfunction, such as hypertension and heart failure, is also a well-documented risk factor for cognitive impairment (Alagiakrishnan et al., 2016; Cannon et al., 2017; Meissner, 2016) and has been associated with changes to brain structure (Kumar et al., 2015; Moon et al., 2018; Suzuki et al., 2017) and function (Bu et al., 2018; Li et al., 2015; Park et al., 2016). As such, both diabetes mellitus and hypertension have been linked to neurological disorders such as stroke (Turin et al., 2016) and dementia (Ninomiya, 2014). One previous report has shown how hypoglycemic conditions in individuals with Type 1 diabetes mellitus results in changes to brain signal MSE (Fabris et al., 2014). We extend these previous findings by showing that the effects of various non-neurological diseases on the brain can be detected by MSE. Together with evidence that MSE changes in response to medical therapies (Farzan et al., 2017; Jaworska et al., 2018; Liang et al., 2014; Okazaki et al., 2015), MSE offers a promising avenue for the development of clinical biomarkers.

## Acknowledgments

This research was supported by a grant from the J. S. McDonnell Foundation to A. R. M.

## CRediT Author Statement

**Kelly Shen**: Conceptualization, Software, Formal Analysis, Validation, Visualization, Writing – Original Draft **Alison McFadden**: Data Curation, Software, Formal Analysis **Anthony R. McIntosh**: Conceptualization, Methodology, Writing – Review & Editing, Supervision, Funding Acquisition

